# NicheScope: Identifying Multicellular Niches and Niche-Regulated Cell States in Spatial Transcriptomics

**DOI:** 10.1101/2025.08.21.671426

**Authors:** Xinyi Yu, Xiaomeng Wan, Leqi Tian, Yuheng Chen, Yuyao Liu, Tianwei Yu, Can Yang, Jiashun Xiao

## Abstract

The functional state of a cell is intrinsically linked to its local microenvironment, or cell niche, a complex milieu formed by multiple interacting cell types. Deciphering how these multicellular niches regulate cell states is fundamental to understanding tissue biology and disease mechanisms, yet remains challenging with current computational approaches. Here, we present NicheScope, a computational framework for transcriptome-wide identification and characterization of Multicellular Niches (MCNs) and their corresponding Niche-Regulated Cell States (NRCSs) from spatial transcriptomics data. NicheScope operates on the principle that a cell’s transcriptional state is associated with its local multicellular composition. It employs a robust statistical approach to jointly model neighborhood composition and transcriptome-wide gene expression, enabling the simultaneous discovery of MCNs, defined by specific combinations of neighboring cell types, and their associated NRCSs, characterized by distinct gene programs. We demonstrate NicheScope’s power and versatility across diverse tissues and platforms, including lymph node, lung adenocarcinoma, and head and neck cancer. NicheScope reproducibly dissected established tissue structures in lymph nodes across tissue regions and platforms, uncovered clinically relevant tumor cell-associated MCNs in lung adenocarcinoma, and revealed shared and condition-specific MCNs in primary and metastatic tumors. Our results establish NicheScope as a powerful, robust, scalable, and interpretable framework for dissecting the spatial and functional organization of complex tissues, providing new insights into multicellular coordination in health and disease.

## Introduction

The state of a cell is determined not only by its intrinsic cell type but also by its interactions within the surrounding microenvironment, or cell niche. Traditionally, these niches have been regarded as static anatomical regions within tissues. However, the advent of spatial transcriptomics (ST) technologies now enables the measurement of gene expression in millions of cells within their native contexts [1, 2, 3, 4]. This capability allows us to reconceptualize cell niches as functional units defined by the interplay of multiple cell types, rather than merely as static structures. Such a paradigm shift presents new opportunities to investigate the complexities of biological processes that are fundamental to tissue organization, development, and disease progression [5, 6, 7].

To formalize this new perspective, we introduce the concepts of the Multicellular Niche (MCN) and the Niche-Regulated Cell State (NRCS). An MCN refers to a specialized microenvironment within a tissue that facilitates the interactions and functions of various cell types, playing a crucial role in regulating cellular states. Cells of the same type can exhibit different states in different MCNs, which we refer to as NRCSs. For instance, in lymph nodes, B cells display different NRCSs across various MCNs: within germinal centers, interactions with follicular dendritic cells (FDCs) and T follicular helper (Tfh) cells promote affinity maturation gene programs, while B cells in the mantle zone exhibit quiescent or early-priming states with distinct functions [8, 9]. A quantitative understanding of MCNs and NRCS is therefore essential for deciphering the complex cellular dialogues that drive tissue homeostasis, immune function, and pathological transformations.

While computational approaches exist to identify cell niches, they often fall short in characterizing Multicellular Niches and Niche-Regulated Cell States. Spatial clustering methods [10, 11, 12, 13, 14], such as CellCharter and scNiche, define cell niches as spatial domains where gene expression patterns are similar or coherent. Although effective for delineating static anatomical regions within tissues, these methods often overlook the complexity of multicellular composition and the functional interactions among various cell types. Ligand-receptor (LR)-based methods [15, 16, 17, 18, 19, 20], such as Giotto and SpaTalk, focus on pairwise signaling analysis using predefined databases, which limits their ability to capture interactions among multiple cell types. As a result, they are not well-suited for characterizing MCNs and NRCSs. Decomposition-based methods, like NCEM, model how the gene expression of a cell is influenced by multiple neighboring cell types. However, these methods often assume that these effects are homogeneous throughout the given tissue, which is a significant limitation. This assumption can hinder the characterization of niche-dependent effects and diminish the ability to effectively characterize NRCSs.

To fully characterize MCNs and NRCSs, we developed NicheScope, built on the principle that the transcriptional state of a cell (the NRCS) is regulated by its surrounding multicellular environment (the MCN), and this relationship can be systematically uncovered by analyzing the association between the cell’s gene expression and the composition of its local neighborhood. To achieve this, NicheScope first quantifies the neighborhood composition for each cell of a target type and identifies genes whose expression is significantly associated with this composition. Then, it uses a multivariate statistical approach that identifies correlated patterns to jointly analyze the selected genes and the neighborhood composition matrix. This allows for the simultaneous detection of MCNs, defined by specific combinations of neighboring cell types, and their corresponding NRCSs, characterized by distinct gene expression programs. By leveraging transcriptome-wide data, NicheScope provides a comprehensive view of niche biology that is not restricted to known ligand-receptor interactions. The framework is computationally efficient, scalable, and supports comparative analysis across multiple conditions.

We demonstrate the power and versatility of NicheScope through its application to diverse tissues and platforms, consistently revealing biologically meaningful MCNs and their corresponding NRCSs. In normal lymph nodes, NicheScope dissected the germinal center by identifying two distinct B cell-associated MCNs: a GC-core niche driving an actively responding NRCS, and a mantle-zone niche supporting a naive-to-priming NRCS. This analysis was reproducible across tissue regions and spatial transcriptomics platforms. Similarly, we further identified a T cell-associated MCN in the T cell zone, where dendritic cells (DCs) guide an NRCS of T cell activation and differentiation. In lung adenocarcinoma, NicheScope uncovered three distinct tumor cell-associated MCNs and indicated clinical relevance: an invasive-front MCN associated with an epithelial-mesenchymal transition (EMT) NRCS and poor survival; a hypoxic MCN driving a pro-angiogenic NRCS; and a bronchial-adjacent MCN linked to a neurodevelopmental NRCS. Further applications revealed an MCN corresponding to tertiary lymphoid structures (TLSs) and resolved stromal heterogeneity into five functionally distinct MCNs. Finally, in a multi-condition analysis of head and neck cancer, NicheScope identified both shared MCNs conserved between primary and metastatic tumors, such as a CAF-driven EMT niche, and condition-specific MCNs reflecting unique tumor-immune dialogues at each site. Together, these results establish NicheScope as a robust and generalizable framework for dissecting the functional organization of complex tissues.

## Results

### Overview of NicheScope method

NicheScope is a unified computational framework designed to systematically identify and characterize Multicellular Niches (MCNs) and their corresponding Niche-Regulated Cell States (NRCSs) from single-cell resolution spatial transcriptomics data. The core principle of NicheScope is that the transcriptional state of a cell is regulated by the multicellular composition of its local microenvironment. The method operationalizes this principle through a two-step statistical approach that integrates spatial and transcriptomic information (Fig. 1).

**Figure 1:**
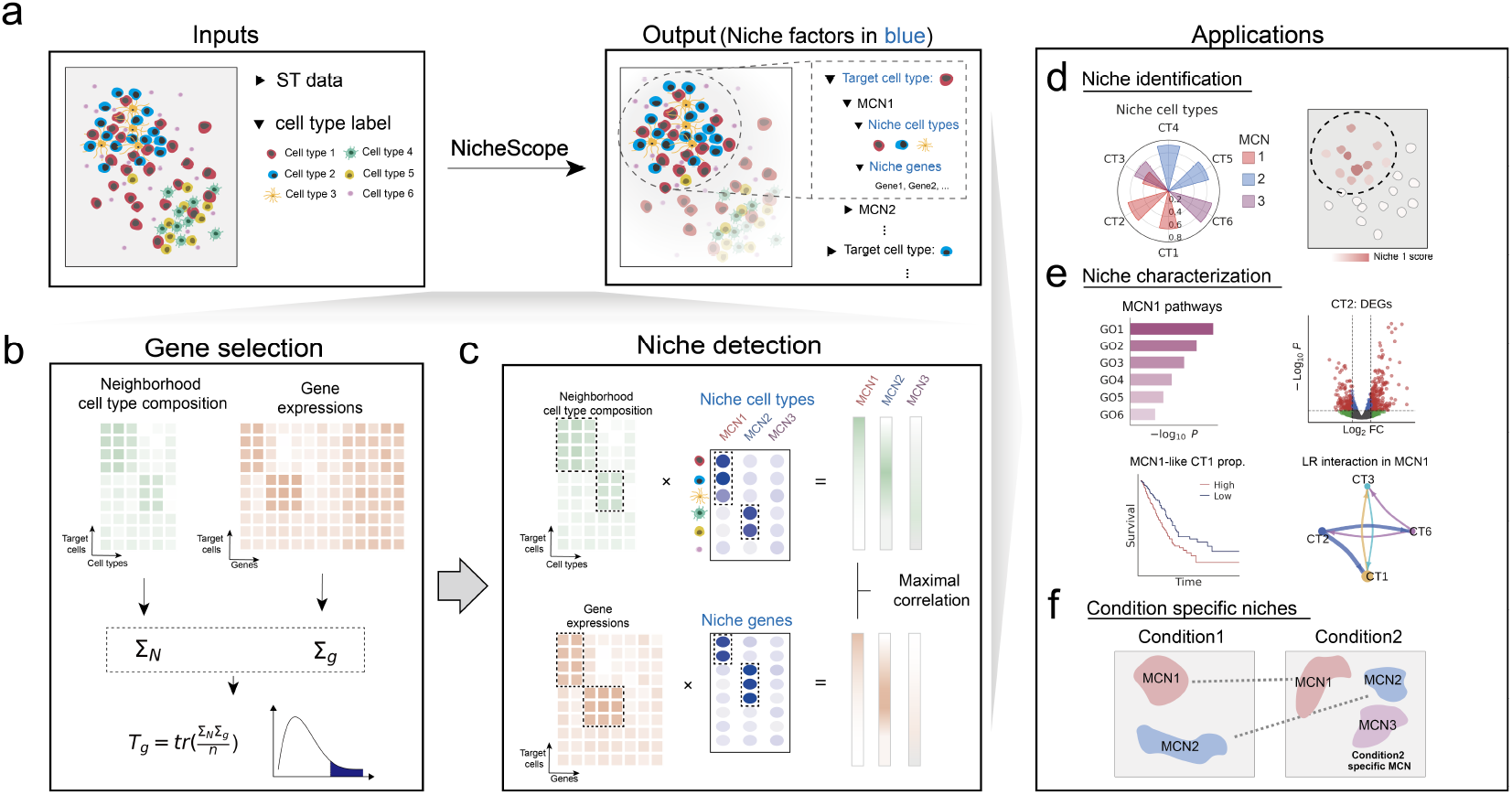
Overview of NicheScope method. **a**-**c**, Given a ST dataset with cell type annotations (**a**), NicheScope focuses on a user-specified target cell type. For each target cell, the neighborhood cell type composition and its gene expression profile are extracted. NicheScope then performs gene-level covariance test to identify genes whose expression patterns are significantly associated with neighborhood composition (**b**). Nonnegative sparse CCA is then applied to jointly model neighborhood composition and candidate gene expression for identifying correlated patterns as successive components (**c**). Each component represents a MCN, defined by niche cell types, and niche genes. These MCNs are the primary output of NicheScope (**a**). **d**, Illustration of multiple detected MCNs and the spatial distribution of an example MCN. **e**, Downstream characterization of one MCN via pathway enrichment, differentially expressed genes of niche cell types, survival associations of MCN-like target cells, and ligand-receptor interactions. **f**, Extension of NicheScope to multi-condition analyses enables discovery of shared and condition-specific MCNs.

First, for each cell of a given target cell type, NicheScope quantifies it local cellular neighborhood by constructing a neighborhood composition matrix where each entry represents the weighted presence of every other cell type in the vicinity, using a Gaussian kernel to prioritize closer cells. This matrix provides a quantitative representation of the spatial context for each target cell. Next, NicheScope identifies genes within the target cell type whose expression patterns are significantly associated with this neighborhood composition. This is achieved through a robust kernel-based covariance test that assesses the statistical dependency between each gene’s expression vector and the neighborhood composition matrix, thereby selecting a set of candidate niche-regulated genes for niche discovery (Fig. 1b).

Finally, to detect the MCNs and corresponding NRCSs, NicheScope applies nonnegative sparse canonical correlation analysis (CCA) to the candidate gene expression matrix and the neighborhood composition matrix. This multivariate technique identifies correlated patterns between the two matrices, yielding a set of components, each representing a distinct MCN (Fig. 1c). For each identified MCN, NicheScope provides two key outputs: (1) a set of “niche cell types” with positive CCA coefficients for neighborhood composition matrix, defining the cellular composition of MCN, and (2) a corresponding set of “niche genes” with positive CCA coefficients for candidate gene expression matrix, characterizing the NRCS of the target cells within that MCN. Nonnegativity ensures interpretability (all contributors act in the same direction), while sparsity highlights the most influential genes and neighboring cell types. Additionally, a “niche score” is calculated for each target cell to quantify its association with the identified MCN, enabling spatial visualization and downstream analysis, including pathway enrichment on niche genes, differential expression of niche cell types inside versus outside the niche, spatially localized ligand-receptor inference within high-scoring regions, and clinical association analyses (e.g., survival analysis) based on MCN-like subpopulations (Fig. 1d,e). The NicheScope framework is also extended to multi-condition analyses, allowing for the joint analysis of multiple datasets or conditions. This extension first identifies shared MCNs that are conserved across conditions and then resolves condition-specific MCNs by analyzing the residual variation unique to each dataset, thus enabling comparative studies of tissue microenvironments (Fig. 1f).

In summary, NicheScope provides a robust approach for dissecting the spatial and functional organization of complex tissues. The entire framework integrates statistical models with efficient algorithms and algebraic optimizations, enabling rapid analysis and low memory usage even for large-scale spatial transcriptomics datasets. For example, a complete niche discovery pipeline for a dataset with 30,000 target cells, 5,000 genes, and 20 cell types can be completed within one minute on a conventional server (Intel i9-14900KF, 24 cores, 128 GiB memory). By offering a formal definition of niches together with a streamlined analysis pipeline, NicheScope delivers a systematic and interpretable characterization of tissue microenvironments, facilitating biological insights across diverse contexts.

### NicheScope reproducibly dissects B cell–associated MCNs in germinal centers of the lymph node

The lymph node’s established structure and well-characterized cell populations make it an ideal benchmark for MCN discovery. We therefore applied NicheScope to a reactive lymph node dataset generated by Xenium (Fig. 2a), from which we extracted two non-overlapping regions (crop 1 and crop 2) for analysis. We used cell2location[21] to annotate cells into 13 major cell types, which contained 34 sub cell types. The spatial distributions of these cell types in the two regions are shown in Fig. S1. We first focused on B cells, a key player in lymph node organization. To gain a spatial overview of B cell arrangement, we clustered B cells based on their neighborhood cell type composition. In crop 1, seven neighborhood composition-based clusters were identified (Fig. 2b). Cluster 2 encompassed multiple follicle-like areas enriched in B cells, FDCs, and T cells, corresponding to GC cores – the primary sites of B cell activation, proliferation, and affinity maturation in reactive lymph nodes [22]. Surrounding these cores, cluster 1 was enriched in memory B (B_mem_) cells and FDCs, representing GC mantle zones. We also observed an isolated follicular structure, cluster 3, resembling a dense GC core without a surrounding mantle zone. Similar clusters localized to GC regions were found in crop 2 (Fig. S3). While this analysis recovered the spatial organization of GC regions, it offered limited insight into their underlying biological functions.

**Figure 2:**
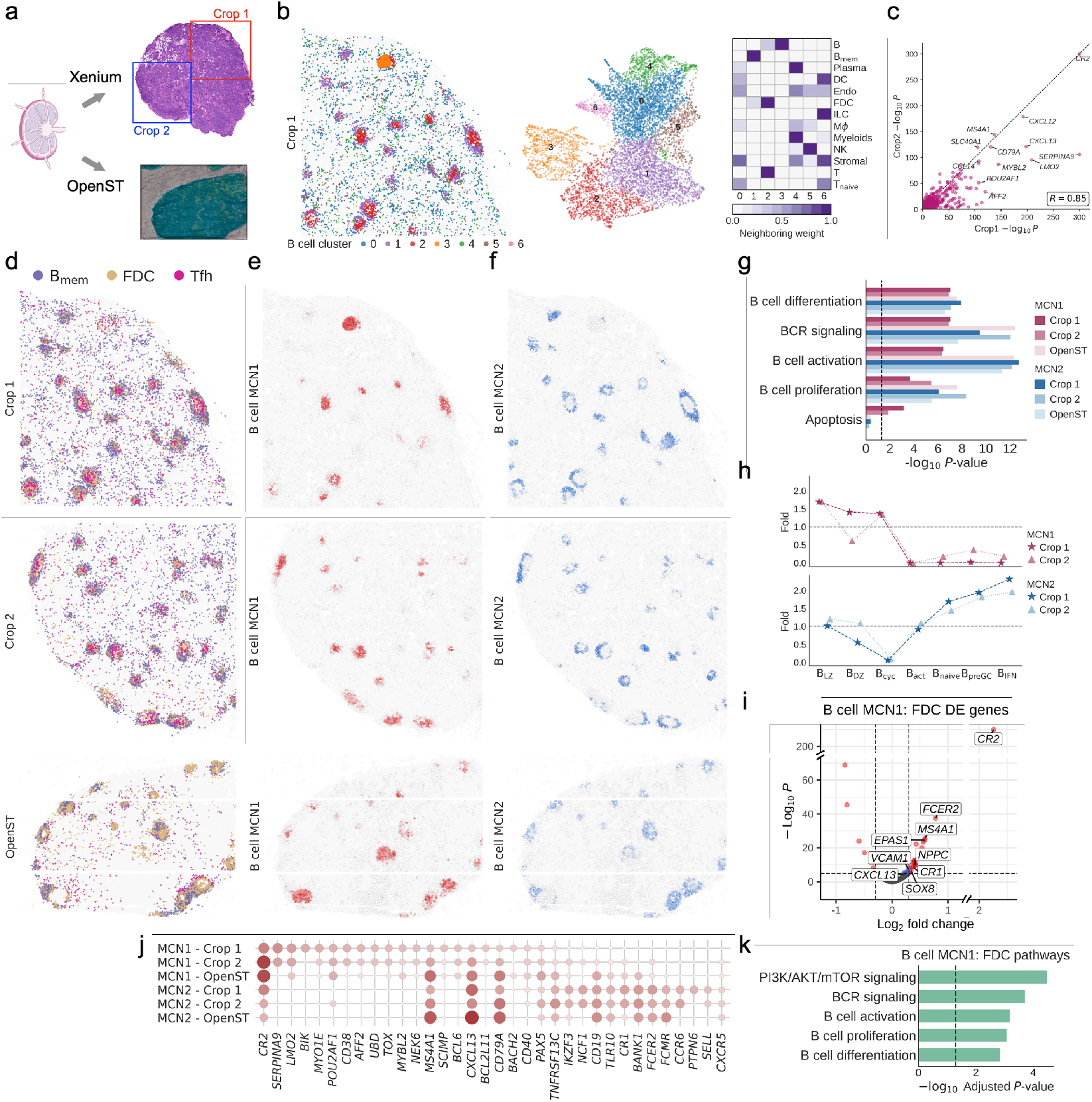
NicheScope reproducibly dissects B cell–associated MCNs in germinal centers of the lymph node. **a**, Overview of datasets used for niche discovery and reproducibility analysis: two non-overlapping regions (crop 1 and crop 2) from Xenium lymph node data and one sample from OpenST lymph node data. **b**, Clustering of B cell neighborhood composition from crop 1 into seven clusters. Shown are their spatial locations, UMAP embedding, and average neighboring cell type weights. **c**, Consistency of gene-level *p*-values from the covariance test between crop 1 and crop 2. **d-f**, Spatial distribution of niche (sub-) cell types – B_mem_ cell, FDC, Tfh cell of B cell-associated MCNs (**d**), niche scores of MCN1 (**e**) and MCN2 (**f**). From top to bottom, each row corresponds to Xenium crop 1, crop 2, and OpenST data, respectively. **g**, Representative enriched pathways for MCN1 and MCN2 across the three datasets. **h**, Enrichment of B cell subtypes within top-scoring (above top 10% niche score) regions for MCN1 and MCN2 in crop 1 and crop 2. **i**,**k**, Differentially expressed genes of FDC within MCN1 (**i**), and representative significantly enriched pathways (adjusted *p*-value *<* 0.05) based on top positively regulated DE genes (logFC *>* 0 and adjusted *p*-value *<* 0.05) in crop 1. **j**, Coefficients of top-ranked niche genes of MCN1 and MCN2 across datasets.

Compared with neighborhood composition analysis, MCN detection using NicheScope not only reveals spatially organized structures of B cells but also uncovers their functional characteristics. To this end, we applied NicheScope to systematically identify MCNs from the perspective of B cells. In the gene selection step, covariance tests were performed separately on crop 1 and crop 2, yielding highly consistent *p*-values across the two crops (Pearson correlation *R* = 0.85) and identifying a large number of significant genes (Fig. 2c). The top 500 genes from each crop were selected as candidates. Using these genes, NicheScope identified two B cell-associated MCNs within GCs and characterized the corresponding NRCS of B cells.

B cell-associated MCN1 corresponded to GC cores, defined by its localization within follicular regions and niche cell types including B cells, FDCs, and T cells (Fig. 2d, Fig. S4,S5). Spatially, MCN1 closely matched the follicle-like neighborhood composition cluster 2 (Fig. 2e). The corresponding NRCS was characterized by a transcriptional program for an active GC response, with niche genes enriched in pathways for receptor (BCR) signaling, activation, proliferation, differentiation, and apoptosis (Fig. 2g,j). For example, niche gene *CR2*, a GC marker which enhances BCR signaling and promotes high-affinity selection [23], was highly expressed in B cells within MCN1 regions (Fig. S6). Notably, NicheScope demonstrated its ability to unify functionally related but structurally distinct regions. In crop 1, an upper region of densely aggregated high-niche-score B cells overlapped with the neighborhood composition cluster 3 (Fig. 2b,d). This area exhibited markedly elevated expression of the GC initiation transcription factor *BCL6* [24] (Fig. S6), together with a scarcity of memory B cells, indicating an earlier-stage GC. Unlike neighborhood composition-based clustering that isolated it as a separate cluster, NicheScope correctly integrated it into the broader GC-core MCN. This highlights how NicheScope moves beyond simple spatial clustering to capture a unified biological process, using niche scores to reflect the relative maturation states within the MCN.

In addition to the transcriptional programs of B cells, the niche cell types within MCN1 further substantiated its GC-core identity. Compared to FDCs outside the niche, differentially expressed genes of FDCs within the niche were enriched in pathways related to GC B cell responses (Fig. 2i,k), including classic GC FDC markers such as *CXCL13, CR2, CR1*, and *VCAM1* [25] (Fig. S8). For instance, *CXCL13* guides the migration of B cells and Tfh cells into GCs [26], while *CR2* and *CR1* facilitate antigen retention and the selection of high-affinity B cells [27]. This illustrates that MCN discovery not only elucidates the NRCS of the target cell type but also enables the inference of transcriptional programs and functional contributions of other constituent cell types.

In parallel, NicheScope identified a second MCN (MCN2) corresponding to the GC mantle zone, forming a distinct ring-like structure around the GC core. MCN2 was characterized by the major niche cell type – memory B cell (Fig. S4,S5), together with enriched B cell subtypes including naive B (B_naive_), pre-GC B (B_preGC_), and interferon-responsive B (B_IFN_) cells (Fig. 2h). This heterogenous cellular composition, comprising undifferentiated B cells that had not yet entered GC cores and differentiated memory B cells that had exited GC cores, suggests MCN2 as a transitional area between the GC core and the T cell zone. While spatially adjacent and sharing foundational pathways like BCR signaling, NicheScope powerfully distinguished MCN1 and MCN2 through their unique NRCSs (Fig. 2g). MCN1 was uniquely enriched for apoptosis signaling and prioritized regulators such as *BCL2L11*, and *BACH2*, indicating that B cells in MCN1 corresponded to an active GC reaction cell state undergoing affinity-based selection and fate commitment [28] (Fig. 2g,j). By contrast, the NRCS of MCN2 prioritized genes of B_naive_ and B_preGC_ cells, including *BANK1*, and *SELL*, reflecting a quiescent or early-priming cell state [29] (Fig. 2j). This difference highlights how distinct NRCSs were captured within the two MCNs. Collectively, NicheScope dissected the spatial and functional continuum of B cell-associated MCNs within GCs, capturing their coordinated yet distinct contributions to the GC response: one focused on B cell proliferation and selection, the other involved in entry, retention, and transitional states. More importantly, these findings were consistently observed across two spatially separated regions – crop 1 and crop 2, demonstrating the cross-region reproducibility of NicheScope in capturing both the spatial architecture and functional specialization of tissue microenvironments.

To assess cross-platform robustness, we further applied NicheScope to an independent OpenST dataset of normal lymph node [4] (Fig. 2a). NicheScope reproducibly resolved two B cell-associated MCNs within GCs, with MCN1 localized to the GC core and MCN2 to the surrounding mantle zone. (Fig. 2d-f). Their niche genes and enriched pathways closely matched those identified in the Xenium data (Fig. 2g,j), demonstrating that NicheScope consistently captured spatially and functionally coherent MCNs across ST platforms.

### NicheScope reveals functionally specialized T cell niches in germinal centers and T cell zones

To further dissect the functional organization of the lymph node, we applied NicheScope to characterize the microenvironments of T cells, which are central to adaptive immunity and exhibit distinct spatial and functional states. We clustered T cells based on their neighboring cell type composition, yielding six clusters in crop 2 (Fig. 3a). Clusters 2 and 4 displayed the most distinct spatial patterns: cluster 2 corresponded to T cells neighboring B cells, FDCs, and memory B cells, indicative of GCs, whereas cluster 4 predominantly neighbored naive T cells, aligning with canonical T cell zones. Similar clusters were observed in crop 1 (Fig. S9). Building on this coarse spatial architecture, NicheScope was then applied to identify MCNs from the perspective of T cells and resolve their corresponding cell states. Covariance tests for candidate gene selection showed highly consistent *p*-values across crops (Pearson *R* = 0.97, Fig. 3b), reflecting robust local context-dependent transcriptional signals. Using these candidates, NicheScope identified two reproducible MCNs of T cell across both crops: one associated with GC regions and the other with T cell zones.

**Figure 3:**
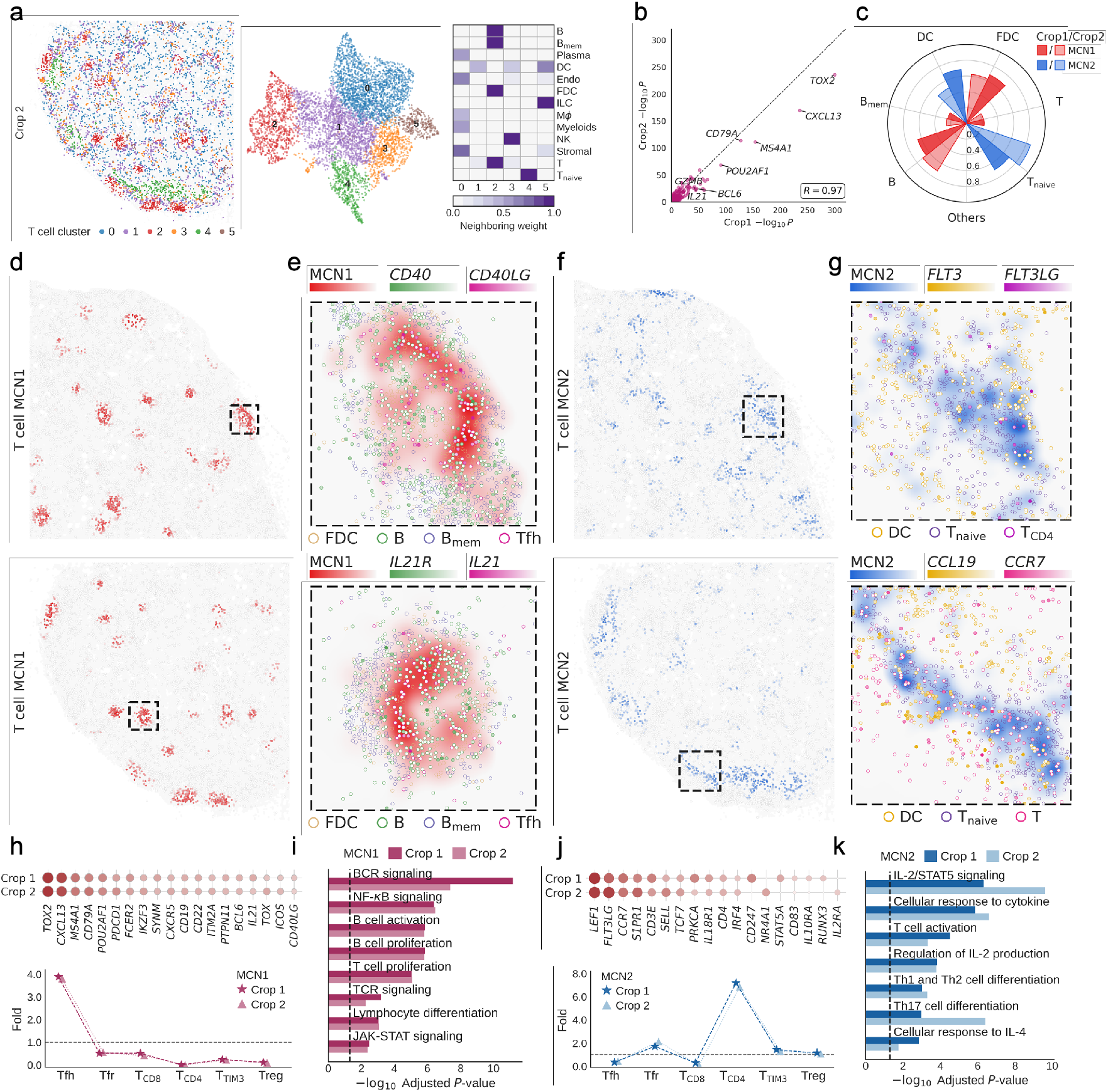
NicheScope reveals functionally specialized T cell niches in germinal centers and T cell zones. **a**, Clustering of T cell neighborhood composition from Xenium crop 2 into six clusters. Shown are their spatial locations, UMAP embedding, and average neighboring cell type weights. **b**, Consistency of gene-level *p*-values from the gene selection step in T cell-associated MCN discovery between crop 1 and crop 2. **c**, Coefficients of major niche cell types (*v* ≥ 0.2) for T cell-associated MCN1 and MCN2 in the two crops. **d**, Spatial distribution of niche score of MCN1 in crop 1 (top) and crop 2 (bottom). **e**, Spatial expression of ligand-receptor pairs related to MCN1: *CD40LG* in Tfh and *CD40* in B cells (top); *IL21* in Tfh and *IL21R* in B cells (bottom); along with niche scores and the distribution of other niche cell types (FDC, B_mem_) in the highlighted region of **d. f**, Spatial distribution of niche score of MCN2 in crop 1 (top) and crop 2 (bottom). **g**, Spatial expression of ligand-receptor pairs related to MCN2: *FLT3LG* in T cells and *FLT3* in DCs (top); *CCR7* in T cells and *CCL19* in DCs (bottom); along with niche scores and the distribution of naive T cells in the highlighted region of **f. h**,**j**, Top: Coefficients of top niche genes of MCN1 (**h**) and MCN2 (**j**) across crop 1 and crop 2. Bottom: Enrichment of T cell subtypes within top-scoring (above top 10% niche score) regions for MCN1 (**h**) and MCN2 (**j**) in crop 1 and crop 2. **i**,**k**, Representative enriched pathways for MCN1 (**i**) and MCN2 (**k**) from crop 1 and crop 2.

T cell-associated MCN1 occupied germinal centers and supported B cell responses, mirroring the spatial distribution of the B cell-associated GC-core MCN identified previously. This finding highlights NicheScope’s ability to capture the similar biological structure from the perspectives of different constituent cell types, thereby providing a multi-faceted view of microenvironment. From the T cell perspective, this MCN defined by the presence of neighboring B cells, FDCs, and was strongly enriched for Tfh, a specialized T subpopulation essential for orchestrating GC reactions (Fig. 3c,d,h, Fig. S10,S11). Although Tfh cells being a minority subpopulation, the identification of this MCN underscores NicheScope’s sensitivity in detecting functionally critical cell states and their corresponding microenvironments. Consistent with its role in supporting GC reaction, MCN1 was functionally enriched in pathways including BCR signaling, B cell activation, T cell receptor (TCR) signaling and T cell proliferation (Fig. 3i).

The niche gene profile of MCN1 provided a detailed functional landscape of the Tfh cell NRCS, with specific genes illuminating their differentiation, migration, effector functions, and interactions with GC B cells (Fig. 3h). First, genes central to Tfh lineage specification and migration were identified, including *BCL6*, the master transcription factor for Tfh commitment and programming [30], and *CXCR5*, the chemokine receptor guiding Tfh cells into B cell follicles [31]. Second, niche genes highlighted Tfh-B cell co-stimulatory interactions, with *ICOS* and *CD40LG* ranked among the top. Specifically, CD154 (encoded by *CD40LG*) expressed on Tfh cells binds CD40 on GC B cells (Fig. 3e, Fig. S12), triggering non-canonical NF-*κ*B signaling and promoting B cell survival, class-switch recombination (CSR), and somatic hypermutation (SHM) [31]. Third, niche genes involved in cytokine signaling further illustrated Tfh effector functions. Notably, *IL21*, encoding the signature cytokine secreted by Tfh cells, was prioritized as a niche gene. IL21 binding to IL21R on B cells activates JAK-STAT signaling, thereby enhancing B cell proliferation and CSR [31] (Fig. 3e, Fig. S12). Finally, *PDCD1*, encoding the hallmark inhibitory receptor PD-1, was also among the top niche genes, reflecting its role in restraining excessive Tfh activity [32]. Together, these niche genes illustrate that NicheScope comprehensively captures the functional cell states of Tfh within the GC-localized MCN.

In parallel, NicheScope resolved a functionally distinct T cell-associated MCN (MCN2) within the T cell zones, capturing the dynamic process of T cell priming and differentiation. MCN2 was mainly composed of T cells, naive T cells, and DCs (Fig. 3c,f, Fig. S10,S11), with CD4^+^ T cells being the most enriched T cell subtype (Fig. 3j). The corresponding NRCS was characterized by pathways for T cell activation, IL2 and IL4 signaling, and Th1/Th2/Th17 differentiation (Fig. 3k), pinpointing MCN2 as a DC-regulated microenvironment supporting T cell activation and lineage commitment.

MCN2 was characterized by a niche gene profile reflecting T cells across successive states (Fig. 3j). Early activation was reflected by TCR signaling genes such as *CD3E, CD247*, and *CD4* [33], as well as genes rapidly upregulated upon TCR stimulation including *IRF4, NR4A1*, and *IL2RA* [34, 35], indicating the presence of antigen-encountering T cells. Furthermore, NicheScope uncovered the molecular basis of the T cell–DC dialogue central to this MCN. Specificaly, *CCR7*, via the *CCR7* -*CCL19* axis, mediates T cell positioning and retention in the T cell zone [36], while *FLT3LG*, secreted by activated CD4^+^ T cells, drives expansion of *FLT3* ^+^ DC precursors, forming a positive feedback loop that sustains T cell priming [37, 38] (Fig. 3g, Fig. S12). As T cells advanced toward differentiation, NicheScope captured the corresponding shift in their transcriptional programs. For example, *RUNX3* and *IL18R1*, pointed to CD8^+^ cytotoxic T cell and Th1 differentiation trajectories [39, 40]. Other niche genes, including *PRKCA, STAT5A*, and *S1PR1* supports clonal expansion[41] and mediate active T cells egress from lymph node [42]. Together, these features underscore MCN2 as a dynamic T cell developmental microenvironment, regulating a spectrum of states spanning naive, activated, differentiating, and migratory T cells.

The T cell-associated MCNs in GCs and T cell zones showed strong consistency across the two non-overlapping tissue crops. Key features – from the selection of candidate genes to niche compositions, spatial distribution, niche genes, and NRCSs – were all reproducibly identified. This high degree of cross-region concordance highlights NicheScope’s ability to robustly resolve MCNs with distinct functional and spatial identities in the lymph node.

### NicheScope uncovers spatially and functionally distinct tumor cell-associated MCNs in lung adenocarcinoma

Lung adenocarcinoma (LUAD) exhibits substantial cellular heterogeneity within its TME, comprising diverse epithelial, immune, and stromal components that collectively shape disease progression and therapeutic response [43, 44]. Spatial transcriptomics technologies enable in situ profiling of these complex microenvironments, offering a powerful opportunity to uncover the spatial organization and cellular composition of tumor-associated MCNs [45]. We applied NicheScope to a Xenium lung adenocarcinoma dataset comprising 21 annotated cell types (Fig. 4a, Fig. S13,S14,S15). Our analyses first centered on tumor cells, treating them as the primary focus for microenvironmental characterization. To gain a preliminary understanding of tumor local context complexity, we clustered tumor cells based on their neighborhood cell type composition. This analysis revealed multiple tumor clusters characterized by distinct immune, stromal, and epithelial surroundings, but their spatial distributions were highly fragmented and lacked coherent structural organization (Fig. 4b). Without incorporating gene expression, the biological relevance of these clusters remained unclear. To further investigate tumor progression, we applied pseudotime inference using slingshot [46], which produced a trajectory that aligned with the expression dynamics of the canonical EMT marker *CDH1*, known to be downregulated during epithelial-mesenchymal transition [47] (Fig. 4c, Fig. S16). Accordingly, the inferred pseudotime was used as a proxy for tumor cell progression in subsequent analyses.

**Figure 4:**
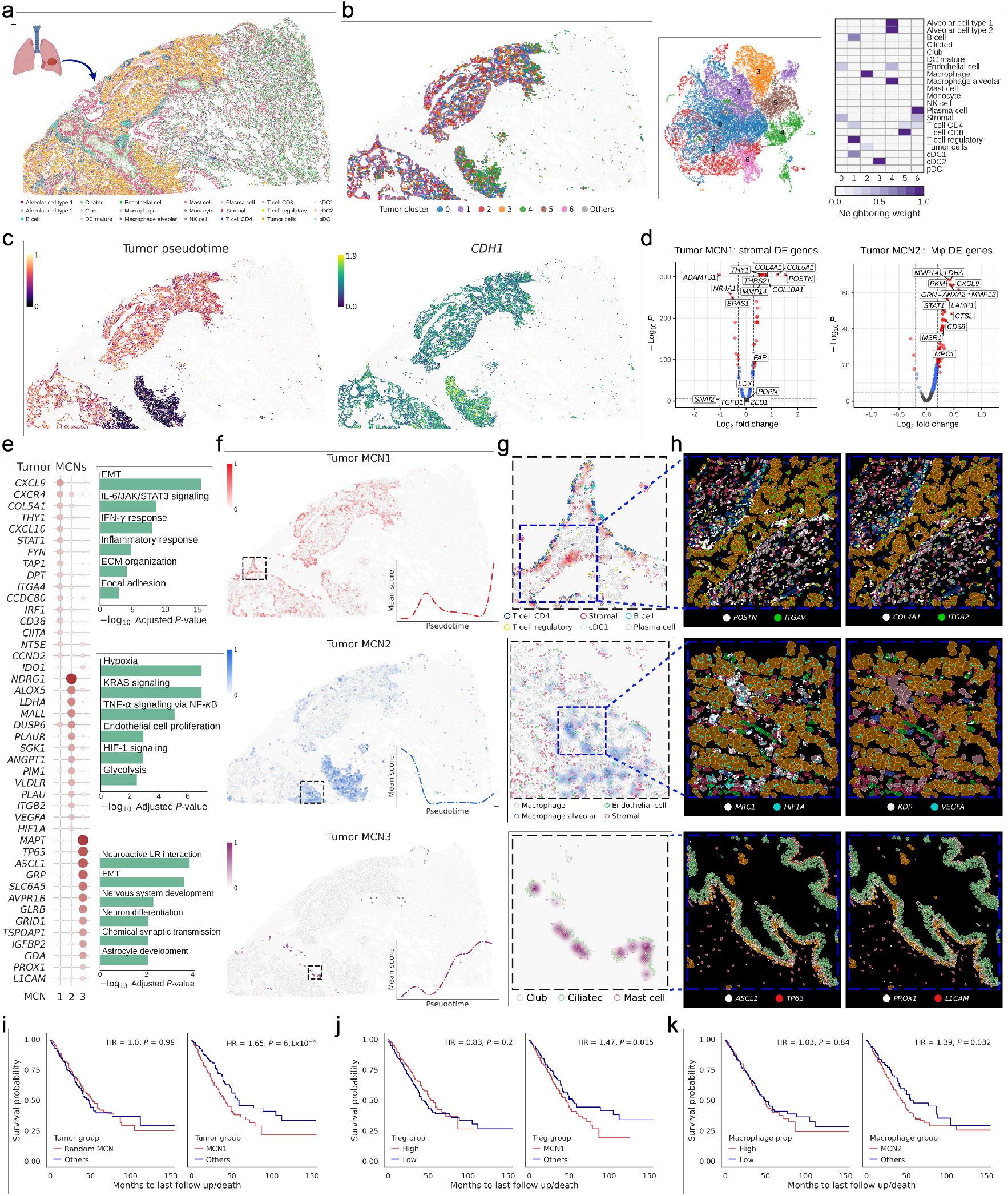
NicheScope uncovers spatially and functionally distinct tumor cell-associated MCNs in lung adenocarcinoma. **a**, Spatial distribution of 21 annotated cell types of the Xenium lung cancer dataset. **b**, Clustering of tumor cell neighborhood composition into seven major clusters. Shown are their spatial locations, UMAP embedding, and average neighboring cell type weights. **c**, Spatial distribution of tumor pseudotime (left) and expression of the EMT marker *CDH1* in tumor cells (right). **d**, Differentially expressed genes of stromal cells within tumor cell-associated MCN1 (left) and macrophages within tumor cell-associated MCN2 (right). **e**, Coefficients of top niche genes (left) and representative enriched pathways (right) of MCN1-3. **f**, Spatial distribution of niche scores and pseudotime of tumor cells within MCN1-3, from top to bottom. **g**, Spatial distributions of niche scores and niche types in the highlighted region of **f. h**, Zoomed-in views of **g** showing cell segmentation and transcript locations of MCN-relevant ligand-receptor pairs or marker genes: *POSTN, ITGAV, COL4A1*, and *ITGA2* for MCN1; *MRC1, HIF1A, KDR*, and *VEGFA* for MCN2; *ASCL1, TP63, PROX1*, and *LICAM* for MCN3. **i**-**k**, Kaplan-Meier survival curves of TCGA-LUAD patients stratified by: a random MCN-like or the relative proportion of MCN1-like tumor cells (**i**), overall stromal cell proportion or the relative proportion of MCN1-like Treg cells (**j**), overall macrophage proportion or the relative proportion of MCN2-like macrophages (**k**).

By applying NicheScope to tumor cells, we systematically resolved the complex tumor microenvironment into three functionally distinct MCNs and their corresponding NRCSs. NicheScope first pinpointed a pro-invasive MCN (MCN1) localized at the peripheral area, a critical region for metastasis due to the presence of lymphatic vessels and capillaries. This MCN was defined by a complex cellular consortium of stromal cells, CD4^+^ T cells, Tregs, B cells, plasma cells, and cDC1s (Fig. 4g, Fig. S17). The corresponding NRCS of tumor cells within this niche was characterized by a transcriptional program enriched for pathways driving EMT, extracellular matrix (ECM) remodeling, and pro-inflammatory signaling (IL6/JAK/STAT3, IFN-*γ*) (Fig. 4e). This functional signature, coupled with the observation that tumor cells in MCN1 were enriched at later pseudotime stages, strongly supports an aggressive, invasive phenotype orchestrated by this specific multicellular microenvironment (Fig. 4f).

NicheScope’s analysis of the immune components within MCN1 revealed a complex, immunosuppressive landscape. By examining the differentially expressed genes of niche cell types, we found that Treg cells within this MCN upregulated canonical immunosuppressive markers (*CTLA4, IL2RG*) and proliferation-associated genes (*CCR4, CCND2*), indicating an actively immunomodulatory Treg population [48, 49] (Fig. S18). NicheScope further implicated a dual-role IFN-*γ* response by identifying its key components (*STAT1, IRF1, CXCL9*) as both niche genes for the tumor cell NRCS and DE genes in Treg and CD4^+^ T cells (Fig. S18). This finding suggests that while the niche promotes T cell recruitment, sustained IFN-*γ* signaling may drive immune tolerance through checkpoint induction such as *IDO1* [50, 51], which also identified as a niche gene (Fig. 4e). Moreover, the identification of *CXCR4* as a niche gene suggests a mechanism where tumor cells exploit immune-derived chemokine gradients for migration and evasion [52].

By characterizing the functional states of niche cell types, NicheScope provided deeper insights into how active stromal remodeling complements this suppressive immune landscape. We first analyzed DE genes of stromal cells within MCN1, and identified the upregulation of multiple CAF markers, such as *POSTN, FAP, LOX*, and *PDPN* [53] (Fig. 4d), confirming a CAF-like transcriptional program known to facilitate EMT [54]. Then, building on this MCN context, spatially-aware ligand-receptor analysis pinpointed key CAF-tumor interactions, including *POSTN* -*ITGAV* and *COL4A1* -*ITGA2*. For instance, stromal-derived *POSTN* binds *ITGAV* on tumor cells to activate FAK/PI3K/AKT signaling, enhancing tumor motility [55], while stromal *COL4A1* engages *ITGA2* on tumor cells to promote adhesion and migration [56]. The spatial co-localization of these ligand-receptor pairs was evident in high-resolution views, where *POSTN* /*COL4A1* -expressing stromal cells were adjacent to *ITGAV* /*ITGA2* -expressing tumor cells (Fig. 4h). This demonstrates how NicheScope not only identifies MCNs but also provides a framework for dissecting the coordinated behaviors of constituent immune and stromal cells that collectively drive an invasive tumor cell state.

Crucially, NicheScope enabled resolving of functionally distinct cell states within MCNs, with direct translation into clinical relevance, as demonstrated by survival analysis of The Cancer Genome Atlas lung adenocarcinoma (TCGA-LUAD) cohort [57]. Patients with a higher abundance of tumor cells transcriptionally resembling the MCN1 associated NRCS exhibited significantly worse overall survival, a correlation that vanished when using a randomly defined tumor cell subpopulation (Fig. 4i, Fig. S20). This prognostic power extended to niche cell types: a high proportion of MCN1-like Tregs was also associated with poorer outcomes, whereas overall Treg abundance showed no such link (Fig. 4j, Fig. S20). These results powerfully demonstrate that NicheScope moves beyond simple cell type enumeration to pinpoint specific, context-dependent cell states that drive patient prognosis. This underscores NicheScope’s potential as a framework for discovering clinically actionable biomarkers by linking spatial microenvironmental architecture to patient outcomes.

NicheScope also resolved a hypoxia-driven MCN (MCN2) that orchestrates early tumor expansion and angiogenesis. This MCN, located adjacent to normal alveolar tissue, was defined by a consortium of macrophages, alveolar macrophages, endothelial cells, and alveolar type 2 cells (Fig. 4f, Fig. S17). The corresponding NRCS revealed a tumor cell state dominated by hypoxia, HIF-1 signaling, glycolysis, and KRAS signaling, consistent with metabolic adaptation and local expansion, a conclusion supported by the enrichment of MCN2-associated tumor cells at earlier pseudotime stages (Fig. 4e,f). A key strength of NicheScope is its ability to characterize the functional states of niche cell types, providing deeper mechanistic insights. By analyzing the DE genes of macrophages within MCN2, NicheScope revealed a shift toward a tumor-supportive, cancer-associated macrophage (CAM) phenotype, marked by the upregulation of genes like *MRC1, MMP14*, and *MSR1* [58] (Fig. 4d). This CAM state was more pronounced than in alveolar macrophages from normal tissue, highlighting a niche-specific polarization (Fig. S19).

NicheScope enabled us to dissect the functional crosstalk driving hypoxia-adapted program. Tumor cells within MCN2 upregulated *HIF1A*, a master regulator of hypoxia response by recruiting and polarizing macrophages [59]. NicheScope captured this interplay, showing that regions with *HIF1A*-high tumor cells were densely populated by M2-like macrophages expressing *MRC1* (Fig. 4h). Simultaneously, NicheScope uncovered a tumor-derived angiogenesis via *VEGFA*, a downstream target of *HIF1A*, was paired with its receptor *KDR* on endothelial cells (Fig. 4h), whose upregulation was confirmed by differential expression analysis within MCN2 (Fig. S19). This demonstrates how NicheScope resolves a complex, pro-angiogenic NRCS sustained by synergistic macrophage and endothelial cell responses.

Survival analysis in the TCGA-LUAD cohort underscored the clinical relevance of this MCN. While overall macrophage abundance showed no prognostic value, a higher proportion of MCN2-like macrophages, as defined by NicheScope, was significantly associated with worse patient survival (Fig. 4k, Fig. S20). This result demonstrates NicheScope’s ability to resolve clinically meaningful, context-dependent cell states that are obscured by simple cell type quantification, directly linking the hypoxia-associated macrophage phenotype to poor prognosis.

Finally, NicheScope uncovered a bronchial-adjacent MCN (MCN3) where tumor cells adopted a neurodevelopmental NRCS linked to an aggressive phenotype. Tumor cells within MCN3 were associated with later pseudotime stages and pathways for EMT, neuron fate specification and neurodevelopmental, suggesting an aggressive phenotype and metastatic potential(Fig. 4e-g, Fig. S17). NicheScope indentified niche genes such as *ASCL1, TP63, PROX1*, and *L1CAM* expressed by tumor cells near bronchial structures (Fig. 4h). The functional relevance of this NRCS of tumor cells is underscored by the roles of these genes in cancer progression: *L1CAM* promotes perineural invasion and directional migration through tumor-nerve interactions [60], while transcription factors like *ASCL1, PROX1*, and *TP63* enhance EMT and plasticity by inducing neural-like or neuroendocrine traits [61, 62]. By resolving this neurodevelopmental NRCS in bronchial microenvironment, NicheScope provides mechanistic clues into neural regulation of LUAD progression.

### NicheScope reveals TLS and stromal heterogeneity in lung adenocarcinoma through MCN detection

Beyond tumor cells, the TME is shaped by diverse immune and stromal populations that strongly influence disease progression and therapeutic responses. To demonstrate the versatility of NicheScope in deciphering complex tissue architecture besides malignant compartments, we further applied it to immune and stromal cells in the same Xenium lung cancer dataset used above.

NicheScope identified a CD4^+^ T cell-associated MCN corresponding to TLSs, functionally resemble secondary lymphoid organs and often localized to tumor margins. TLSs have been reported to be associated with enhanced anti-tumor immune responses and favorable prognosis in lung cancer [63]. NicheScope precisely captured these structures, where the identfied MCN spatially co-localized with regions exhibiting high TLS signature scores at the tumor-stromal interfaces (Fig. 5a-c). This MCN was defined by hallmark features of TLSs, including a canonical cellular composition (CD4^+^ and CD8^+^ T cells, Tregs, B cells, and DCs) and pathways of lymphocyte activation, differentiation, and chemokine signaling (Fig. 5c). Zoomed-in views revealed the identified TLS architecture, with stromal cells forming an outer scaffold surrounding an inner zone enriched in T cells, B cells, and DCs (Fig. 5d). Critically, NicheScope resolved the key molecular interactions driving this structure, highlighting active chemokine axes such as *CXCL13* -*CXCR5* and *CCL19* -*CCR7* responsible for recruitment and organization of lymphocytes [64] (Fig. 5d, Fig. S21). This demonstrates NicheScope’s ability to decouple complex immune structures by integrating their cellular, functional, and architectural features into a single coherent MCN.

**Figure 5:**
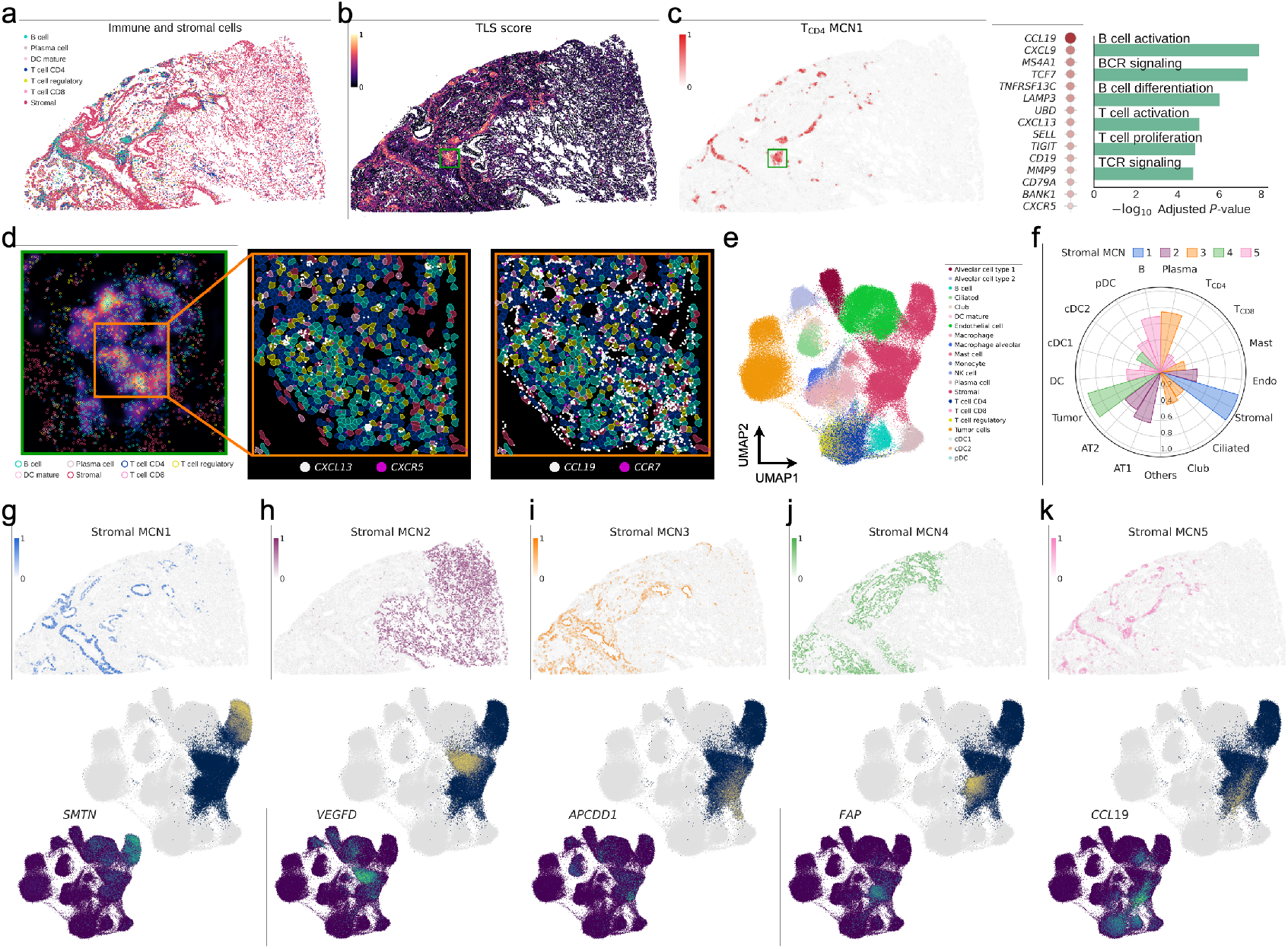
NicheScope reveals TLS and stromal heterogeneity in lung adenocarcinoma through MCN detection. **a**, Spatial distribution of immune and stromal cells in the Xenium lung cancer dataset. **b**, Spatial distribution of TLS signature score. **c**, Spatial distribution of niche score (left), coefficients of top niche genes (middle) and representative enriched pathways (right) of CD4^+^ T cell-associated MCN1. **d**, Left: Spatial distribution of TLS scores and immune and stromal cells in the highlighted region of **b**. Zoomed-in views showing cell segmentation and transcript locations of TLS-associated ligand-receptor pairs: *CXCL13* -*CXCR5* (middle) and *CCL19* -*CCR* (right). **e**, UMAP embedding of 21 cell types. **f**, Coefficients of major niche cell types (*v* ≥ 0.2) for stromal cell-associated MCN1-5. **g**-**k**, Spatial distribution of niche scores, UMAP embeddings of niche scores and representative niche genes of stromal cell-associated MCNs, from left to right.

Stromal cells are essential to tissue architecture and immune regulation in the TME. In Xenium lung cancer dataset, they were annotated as a single, seemingly homogeneous population (Fig. 5e), yet this category likely encompassed diverse subtypes or cell states. Applying NicheScope to stromal cells, we successfully resolved this apparent uniformity by dissecting the stromal compartment into five functionally and spatially distinct MCNs, each corresponding to a unique stromal cell state (Fig. 5f, Fig. S22). MCN1 was located around bronchi and marked by the niche gene *SMTN* encoding smoothelin [65], indicating a bronchial smooth muscle cell state (Fig. 5g). MCN2 was distributed across normal alveolar regions enriched in fibroblasts supporting alveolar structure, regeneration, and immune homeostasis, as evidenced by the niche gene *VEGFD* involved in lymphangiogenesis is selectively expressed in this MCN [66] (Fig. 5h). MCN3 resided in the interstitial zone between bronchi and alveoli, and the expression of niche genes *APCDD1* and *PDGFRA* indicates a interstitial fibroblasts cell state involved in lung development and repair [67] (Fig. 5i). MCN4 was confined to the tumor region and exhibited transcriptional features of CAFs, marked by exclusive upregulation of *FAP* [68] (Fig. 5j). MCN5 spatially overlapped with TLSs and showed high expression of *CCL19*, consistent with its role in supporting TLS organization and immune cell recruitment [69] (Fig. 5d,k). This comprehensive dissection demonstrates NicheScope’s ability to deconstruct a complex cell population into its constituent functional states, revealing a hidden layer of stromal heterogeneity critical to tissue organization.

### Multi-condition niche discovery using NicheScope: a case study in primary and metastasis HNSCC

To showcase NicheScope’s power in dissecting microenvironmental evolution, we applied its multi-condition framework to a matched pair of OpenST datasets from a head and neck squamous cell carcinoma (HNSCC) patient, capturing the primary tumor and its lymph node metastasis [4] (Fig. 6a). Primary and metastatic tumors exhibit conserved and divergent microenvironmental features: primary tumors often maintain organized epithelial-immune architectures, whereas metastatic sites show reduced epithelial differentiation and greater immune evasion [70]. In line with this, neighborhood enrichment analysis revealed both shared and site-specific spatial relationships among cell types (Fig. S23), motivating a joint analysis to identify MCNs shared across sites or unique to tumor progression. We first examined tumor cell neighborhoods in each dataset, identifying seven major clusters: five shared across sites, e.g., CAF-enriched (cluster 2), keratin pearl (cluster 3), CAM-enriched (cluster 4); one primary-specific (DC–endothelial, cluster 5), and one metastasis-specific (B cell–enriched, cluster 6) (Fig. 6b). These patterns highlighted both shared and site-specific tumor architectures, but their functional implications were lacking. We therefore leveraged NicheScope to systematically deconstruct this landscape, aiming to identify tumor cell-associated MCNs and NRCSs that are either conserved across both sites or emerge as unique hallmarks of primary versus metastatic disease (Fig. 6c).

**Figure 6:**
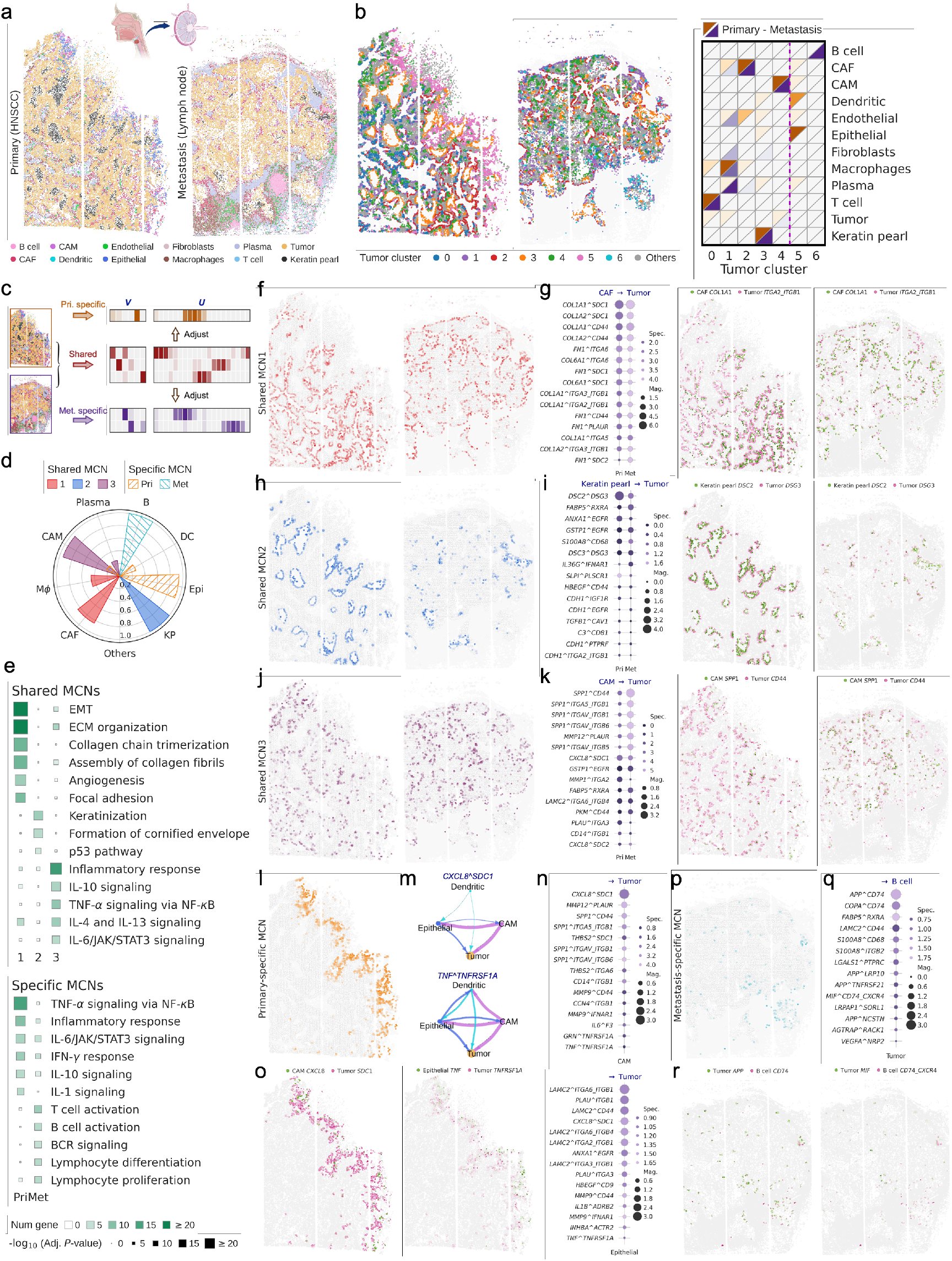
Multi-condition niche discovery using NicheScope: a case study in primary and metastasis HNSCC. **a**, Spatial distribution of cell types in HNSCC primary tumor and lymph node metastatic tumor. **b**, Clustering of tumor cell neighborhood composition into seven clusters per dataset. Shown are spatial locations and average neighboring cell type weights. **c**, Overview of NicheScope for identifying shared and condition-specific tumor cell-associated MCNs across datasets. **d**, Coefficients of major niche cell types (*v* ≥ 0.2) for shared MCN1-3, the primary- and metastasis-specific MCNs. **e**, Representative enriched pathways of shared (top) and condition-specific (bottom) tumor cell niches. **f**,**h**,**j**, Spatial distribution of niche score of shared MCN1 (**f**), MCN2 (**h**), and MCN3 (**j**) in both tumors. **g**,**i**,**k**, Left: magnitude and specificity of ligand-receptor interactions in shared MCN1-3: CAFs to tumor cells (**g**), keratin pearl tumor cells to tumor cells (**i**), CAMs to tumor cells (**k**). Right: spatial expression of representative ligand-receptor pairs-*COL1A1* -*ITGA2 ITGB1* (**g**), *DSC2* -*DSG3* (**i**), and *SPP1* -*CD44* (**k**). **l**, Spatial distribution of the primary-sepcific tumor cell niche score. **m**, Magnitude of *CXCL8* -*SDC1* (top) and *TNF* -*TNFRSF1A* (bottom) among tumor cells and primary-specific tumor cell types (epithelial cell, CAM and DC). **n**, Magnitude and specificity of top ligand-receptor interactions from CAMs (top) and epithelial cells (right) to tumor cells in the primary-specific MCN. **o**, Spatial expression of *CXCL8* -*SDC1* (left) and *TNF* - *TNFRSF1A* (right) in corresponding sender-receiver cell types. **p**, Spatial distribution of the metastatic-sepcific tumor cell niche score. **q**, Magnitude and specificity of top ligand-receptor interactions from B cells to tumor cells in the metastasis-specific MCN. **r**, Spatial expression of representative ligand-receptor pairs in the metastasis-specific MCN: *APP* -*CD74* (left) and *MIF* -*CD74* /*CXCR4* (right).

NicheScope identified three major shared tumor cell-associated MCNs. Shared MCN1 represented a fibroblast-rich microenvironment that orchestrated an EMT and ECM remodeling NRCS in tumor cells (Fig. 6d,f). Pathway enrichment confirmed its involvement in ECM organization and collagen formation (Fig. 6e, Fig. S24). NicheScope further resolved the underlying molecular dialogue, revealing extensive ECM-tumor crosstalk where CAF-derived collagens and fibronectins engaging integrin receptors on tumor cells. For example, *COL1A1* - *ITGA2 ITGB1* interaction was consistently detected in both primary and metastatic tumors (Fig. 6g, Fig. S26), which is known to drive ECM remodeling and invasion [71]. These results indicate that tumor cells within shared MCN1 adopted a conserved NRCS characterized by active interaction with fibroblasts that promote invasion.

A keratinization-focused MCN (shared MCN2) was identified at the periphery of keratin pearls, a defining feature of well-differentiated HNSCC (Fig. 6d,f). The corresponding NRCS was defined by pathways of keratinization and cornified envelope formation (Fig. 6e, Fig. S24) and a suite of niche genes including *LCE3E* and *KRT17* [72, 73]. NicheScope pinpointed the *DSC2* -*DSG3* interaction, involving desmosomal cadherins crucial for intercellular adhesion [74], as the dominant molecular axis maintaining this structure. The elevated expression of *DSG3* further underscored a well-differentiated tumor cell state [75] (Fig. 6i, Fig. S27).

NicheScope further resolved a conserved inflammatory niche (shared MCN3), revealing a CAM-driven microenvironment scattered throughout the tumor parenchyma (Fig. 6d,j). The corresponding NRCS was defined by a tumor cell response to cytokine-mediated and TNF-*α*/NF-*κ*B signaling (Fig. 6e, Fig. S24). LR analysis highlighted interactions involving CAM signature genes such as *SPP1, MMP12*, and *CXCL8* [76, 77], with *SPP1* -*CD44* as a key pair promoting tumor adhesion, migration, and immunosuppressive signaling [78, 79]; this interaction was observed in both datasets and exhibited stronger co-expression in the metastasis (Fig. 6k, Fig. S28).

Beyond conserved tumor microenvironment, NicheScope also uncovered condition-specific MCNs, highlighting unique tumor-immune dialogues at each disease stage. The primary-specific tumor cell-associated MCN localized in a distinct epithelial-immune interface enriched in epithelial cells, CAMs and DCs (Fig. 6d,l). The corresponding NRCS was characterized by a complex inflammatory programs, including TNF-*α* signaling via NF-*κ*B, and cytokine-mediated signaling, which partially overlapping with shared MCN3 but unique to the primary tumor (Fig. 6e, Fig. S25). Within primary-specific MCN, NicheScope uncovered multiple LR interactions between tumor cells and niche cell types that mediate inflammatory responses, proliferation, and ECM remodeling (Fig. 6n). For example, *CXCL8* -*SDC1*, known to promote inflammation, angiogenesis, and migration [80, 81], showed varying co-expression across several interacting cell type pairs (Fig. 6m,o, Fig. S29). Another LR interaction, *TNF* -*TNFRSF1A*, as a key axis of TNF-*α* signaling, was broadly co-expressed among diverse cell types (Fig. 6m,o, Fig. S29), exerting dual effects by inducing NF-*κ*B–mediated cytokines (e.g., IL6, IL8) while contributing to immunosuppression and tumor progression [82]. These features delineate an NRCS of tumor cells driven by epithelial–immune interactions and marked by both pro-inflammatory and immunosuppressive signaling.

In the metastatic lesion, NicheScope uncovered a unique MCN at the tumor-immune interface, defined by a B cell-rich microenvironment adjacent to TLS-like structures (Fig. 6d,p). Metastasis-specific MCN was enriched for pathways related to B/T cell activation and lymphocyte proliferation and differentiation (Fig. 6e, Fig. S25), reflecting a distinct immune microenvironment adjacent to TLS-like regions. Several interaction pairs known to function in the tumor–immune interface were identified in this MCN, including *MIF* -*CD74* and *MIF* -*CXCR4*, which mediate inflammatory signaling, immune evasion, and regulation of B cell activity [83, 84] (Fig. 6r,p, Fig. S30). Beyond these well-established interactions, our analysis also revealed prominent underappreciated interactions between tumor cell-derived ligands *APP* /*COPA* and receptor *CD74* on B cells (Fig. 6r,p, Fig. S30). While direct evidence for such signaling is lacking, recent studies have implicated these crosstalk in immunosuppression and poor prognosis in cervical, pancreatic, and testicular cancers [85, 86, 87, 88]. Our observations demonstrate that tumor cells in the metastasis-specific MCN adopt an NRCS shaped by B cell–mediated immune signaling, and offer new hypotheses for future investigation into tumor–B cell communication and tumor immune evasion mechanisms.

In summary, the analysis of primary and metastatic tumors highlights the capability of NicheScope to uncover both shared and condition-specific MCNs and corresponding NRCSs across distinct datasets. The insights gained from these identified MCNs provide a strong foundation for future investigations comparing multicellular microenvironments across patients, tumor stages, and therapeutic responses.

## Discussion

Dissecting the spatial organization of multicellular niches and their corresponding niche-regulated cell states is fundamental for understanding tissue function and disease mechanisms. While existing methods for niche discovery have provided valuable insights, they often face limitations. Spatial clustering approaches can delineate anatomical regions but may overlook the functional impact of multicellular composition. Ligand-receptor-based methods are constrained to known pairwise interactions, limiting their ability to capture the complexity of MCNs. Decomposition-based methods often assume homogeneous effects across a tissue, which can obscure the diversity of MCNs and NRCSs. NicheScope addresses these limitations by systematically modeling the association between a cell’s transcriptomic profile and its local multicellular neighborhood, thereby enabling unbiased detection and characterization of MCNs defined by multiple interacting cell types and their associated NRCSs in spatially localized regions. Through comprehensive analysis across diverse tissue types and spatial transcriptomics platforms, NicheScope demonstrates robust and reproducible performance, revealing both shared and condition-specific cell niches and uncovering novel microenvironments linked to clinical outcomes. In summary, NicheScope provides a powerful and interpretable framework for elucidating the multicellular architecture of complex tissues, shedding light on how spatially organized cellular coordination shapes tissue function and pathology.

Despite these strengths, NicheScope has several limitations. First, it is primarily designed for single-cell resolution spatial transcriptomics data. For lower-resolution platforms such as 10x Visium[89], where each spot may contain multiple cells, accurate dissection of cell niches becomes more challenging and may require additional deconvolution steps or integration with single-cell data [90]. Nevertheless, with the increasing availability and adoption of commercial platforms offering single-cell or near-single-cell resolution (e.g., Xenium, Visium HD [91]), NicheScope is well positioned to leverage these technological advances and facilitate high-resolution spatial data mining. Second, the accuracy of NicheScope relies on precise cell type annotation, as the model explicitly links gene expression profiles to neighboring cell types. While annotation of major cell types is generally feasible, distinguishing fine-grained subtypes remains difficult due to transcriptional similarity and technical noise [92]. To mitigate potential false positives, we recommend performing niche identification using major cell types initially when the fine-grained subtypes are not well annotated with high confidence. This approach allows for the identification of biologically meaningful niches at a higher level of granularity, followed by enrichment analysis of subtypes within identified niches. For example, to enhance robustness and interpretability, our analysis of B/T cell niches in the lymph node was conducted using 13 major cell types instead of 34 fine-grained subtypes.

Additional limitations include the modeling assumptions and spatial context definition in NicheScope. Our method captures linear dependencies underlying cell-cell interactions in spatial transcriptomic data. While linear models have demonstrated strong performance in related tasks [93], they may not fully capture complex, nonlinear interactions present in some biological systems. Nevertheless, the interpretability and computational efficiency of linear models make them attractive for large-scale spatial analyses. Besides, NicheScope requires the specification of a Gaussian kernel bandwidth to define the spatial context of each cell. In fact, our analysis indicate that NicheScope is robust to bandwidth selection across a range of values. Alternatively, the method supports an adaptive strategy based on the Cauchy *p*-value combination rule [94], enabling gene-specific bandwidths and allowing simultaneous modeling of both short- and long-range cellular interactions (see Methods for details).

Looking forward, NicheScope offers several promising avenues for extension. Integrating multi-omics data, such as spatial proteomics [95] or epigenomics [96], could provide a more comprehensive view of cell niche composition and function. Addressing batch effects and harmonizing data from different spatial transcriptomics platforms will further enhance the generalizability and applicability of NicheScope in large-scale, multi-center studies. Moreover, extending the framework to accommodate spatial-temporal data [97] would enable the investigation of dynamic changes in cell niches during development, disease progression, or therapeutic intervention. Collectively, these future directions will further empower NicheScope to advance our understanding of tissue organization and multicellular interactions across diverse physiological and pathological conditions.

## Methods

### NicheScope under a single condition

To detect and characterize MCNs and their associated NRCSs from spatial transcriptomics data, NicheScope employs a unified statistical framework that leverages both transcriptional and spatial information. The core idea of NicheScope is that the transcriptional states of target cells are shaped by the cellular composition of their surrounding microenvironment. To operationalize this idea, NicheScope proceeds in two main steps: (i) **gene selection**, which identifies candidate signature genes of MCNs by detecting genes in the target cell type whose expression patterns are strongly associated with neighborhood cell type composition; and (ii) **niche detection**, which uses nonnegative sparse CCA to jointly infer MCNs and the corresponding NRCSs.

Consider a 2-dimensional single-cell resolution spatial transcriptomics dataset comprising *n* cells and *q* genes in the set 𝒢. The dataset contains a gene expression matrix **X** ∈ ℝ^*n×p*^ and the spatial coordinates of cells **Z** ∈ ℝ^*n×*2^ within the tissue. Suppose there are *m* cell types, and each cell has been annotated with a cell type label. Let ***y*** ∈ {1, 2, …, *m*}^*n*^ denote the cell type annotation vector. Focusing on a target cell type *c*, we construct a **neighborhood cell type composition matrix** 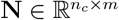, where *n*_*c*_ denotes the total number of cells labeled as cell type *c*. This matrix characterizes the local cellular context of each target cell. The (*i, k*)-th element of **N**, representing the weighted presence of cell type *k* in the neighborhood of the *i*-th target cell, is defined as follows:

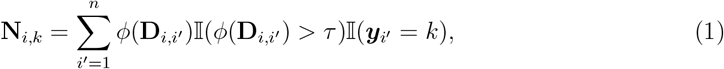

where 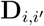 denotes the Euclidean distance between *i* and 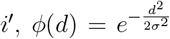 is the Gaussian kernel with bandwidth *σ*. The threshold *τ* defines a minimum kernel value to filter out distant interactions. 𝕀(·) denotes the indicator function. This construction of the cell type composition matrix incorporates distance-dependent weighting, reflecting the principle that proximal cells have a stronger influence on the target cell.

### Gene selection

The selection of candidate signature genes of MCNs is based on the fundamental assumption that such genes exhibit expression patterns associated with the neighborhood composition of the target cell type. In contrast, genes that are uniformly expressed across the tissue microenvironment are not of primary interest. To identify genes that meet this criterion, we assess the statistical association between each gene’s expression and the neighborhood cell type composition matrix. For a given gene *g*, let 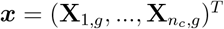 denote its expression vector across all target cells. Intuitively, if ***x*** is correlated with **N**, target cells with similar neighborhood cell type compositions should exhibit similar expression levels of gene *g*. This relationship can be quantified by comparing the covariance structures of ***x*** and **N**. After zero-centering both variables, we define their covariance matrices as:

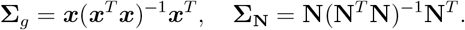

Under the null hypothesis that ***x*** and **N** are statistically independent, the test statistic *T*_*g*_

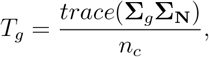

has the same asymptotic distribution as a linear combination of *χ*^2^ random variables [98]:

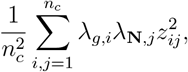

where *λ*_*g,i*_, *λ*_**N**,*j*_, *i, j* = 1, …, *n*_*c*_ are the eigenvalues of Σ_*g*_ and Σ_**N**_, respectively. The *p*-value of the covariance test can be calculated through Hall-Buckley-Eagleson’ approximation method [99, 100]. After calculating *p*-values of all the genes and adjust for multiple testing, we obtain a candidate gene set 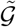 consisting of *q*^*′*^ genes highly correlated with neighborhood cell type composition. The corresponding gene expression matrix is denoted as 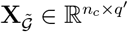.

To ensure computational efficiency and manage memory requirements for large spatial datasets, following [101], we employ several algebraic techniques to avoid computing and operating on the *n*_*c*_ *× n*_*c*_ covariance matrices **Σ**_*g*_ and **Σ**_**N**_ directly. First, the eigenvalues of **Σ**_*g*_ and **Σ**_**N**_ are equivalent to the eigenvalues of (***x***^*T*^ ***x***)^−1^***x***^*T*^ ***x*** and (**N**^*T*^ **N**)^−1^**N**^*T*^ **N**, where the former is a scalar and the latter is a *m × m* matrix that only need to be computed once. Second, we can express the test statistic as:

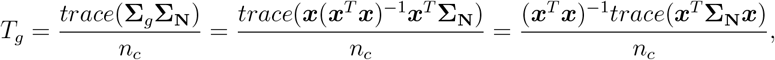

where the test statistic can be expressed in terms of ***x***^*T*^ ***x*, N**^*T*^ **N**, and **x**^*T*^ ***N***. These calculations scale linearly with the number of target cells *n*_*c*_, resulting in a significant reduction in computational complexity and memory footprint compared to operations involving *n*_*c*_ *× n*_*c*_ matrices.

While a single Gaussian kernel bandwidth *σ* is typically used to construct the neighborhood composition matrix, the covariance test framework can be naturally extended to incorporate multiple spatial scales. Specifically, for each gene, one may perform the covariance test across a range of bandwidth values, resulting in a set of *p*-values corresponding to different spatial scales. These individual *p*-values can be aggregated using the Cauchy combination rule [94], a method designed to enhance statistical power by integrating signals across multiple tests. This extension provides an alternative strategy for selecting candidate genes that may exhibit spatial associations at varying scales. In our study, we used a fixed bandwidth for gene selection, but the multi-scale approach remains a useful option when aiming to capture a broader set of candidate genes.

### Niche detection

Matrices 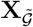 and **N** represent the candidate gene expression profiles and the corresponding neighborhood composition information for each target cell, respectively. MCNs are identified by investigating their associations through nonnegative sparse CCA, which iteratively computes canonical coefficient vectors maximizing the correlation between the two variable sets under nonnegativity and sparsity constraints. Specifically, let **X**^(1)^ and **N**^(1)^ denote the standardized versions of 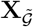 and **N**. With respective cell type *c*, the first MCN, referred to as MCN1, is defined as the first nonnegative sparse CCA component, obtained by solving the following optimization problem [102]:

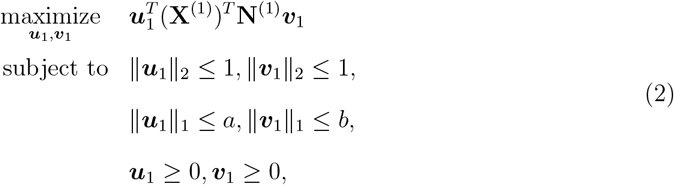

where 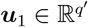 and ***v***_1_ ∈ ℝ^*m*^ are the **gene coefficients** and **cell type coefficients**, respectively. The nonnegative constraints are imposed because allowing both positive and negative values in ***u***_1_ and ***v***_1_ would complicate interpretation of the MCN. The sparsity parameters *a, b* ∈ [0, 1] control how many genes and cell types are included to characterize the MCN. These parameters are user-defined to balance interpretability and inclusiveness, depending on the biological context. The sparsity constraints are imposed to prioritize the cell types and genes that contribute most to this MCN. We define **niche genes** as genes with positive loadings in ***u***_1_, i.e., {*g*: ***u***_1,*g*_ *>* 0}, and **niche cell types** as cell types with positive loadings in ***v***_1_, i.e., {*k*: ***v***_1,*k*_ *>* 0}. Together, the niche cell types and niche genes provide the primary descriptors of the MCN, with the magnitude of each CCA coefficient reflecting its relative contribution. Notably, the niche genes specifically define the corresponding NRCS. The linear combinations **X**^(1)^***u***_1_ and **N**^(1)^***v***_1_ are referred to as the niche gene score and niche composition score, respectively. Their elementwise product defines the **niche score**, given by ***s***^(1)^ = (**X**^(1)^***u***_1_) ⊙ (**N**^(1)^***v***_1_), which quantifies how strongly each target cell belongs to MCN1. Target cells with high niche scores are considered to belong to MCN1 and exhibit the corresponding NRCS of this niche.

To obtain subsequent MCNs, we construct adjusted input matrices that account for the contribution of the previous component. For the second MCN, referred to as MCN2, the adjusted matrices are given by

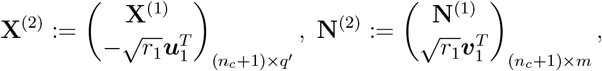

where 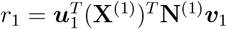. By solving the optimization problem in Eq. (2) on these adjusted matrices, we obtain the second nonnegative sparse CCA component, which defines MCN2. The resulting coefficient vectors, ***u***_2_ and ***v***_2_, define the niche genes and niche cell types of niche 2. Analogous to niche 1, niche 2 scores can be calculated, and target cells with high niche 2 scores display the corresponding NRCS of MCN2. Repeating the above procedure *L* times yields *L* MCNs in total. Of note, unlike the classical CCA, the resulting *L* sets of niche gene scores and niche composition scores are not guaranteed to be orthogonal, due to the nonnegativity and sparsity constraints.

### NicheScope under multiple conditions

NicheScope can be extended to multi-condition niche identification, enabling the detection of both shared and condition-specific MCNs across multiple spatial transcriptomics datasets. For example, it can be applied to identify common and unique MCNs from spatial transcriptomic data of primary and metastatic tumor samples. Below, we describe the method in the case of two datasets. The extension to three or more conditions follows naturally.

Consider two 2-dimensional spatial transcriptomics datasets with *n*_1_ and *n*_2_ cells, respectively. Let 𝒢_1_ and 𝒢_2_ denote the gene sets in the two datasets, with sizes *q*_1_ and *q*_2_. Define 𝒢_0_ = 𝒢_1_ ∩ 𝒢_2_ as the set of shared genes with size *q*_0_, and 𝒢 = 𝒢_1_ ∪ *𝒢*_2_ as the union of all genes with size *q*. The corresponding gene expression matrices are denoted as 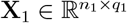 and 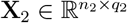. Assume that all the cells have been annotated with cell types. Let 𝒞_1_ and 𝒞_2_ be the cell type sets in the two datasets, with sizes *m*_1_ and *m*_2_, respectively. The shared cell type set is 𝒞_0_ = 𝒞_1_ ∩ *𝒞*_2_ of size *m*_0_, and the union of all cell types is 𝒞 = 𝒞_1_ ∪ *𝒞*_2_ of size *m*. In general, the goal is to compare tissue sections with substantial biological similarity, such as the same tissue type from different individuals, or a tissue under different conditions (e.g., normal vs. disease). Under this setting, most genes and cell types are expected to be shared across the datasets, although a limited number of dataset-specific genes or cell types may also be present.

Let a shared cell type *c* be the target cell type, and denote the number of target cells in the two datasets as 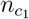 and 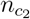, respectively. Based on Eq. (1), we compute the neighborhood cell type composition matrices 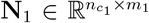 and 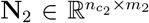 for cell type *c*. As in the single-condition NicheScope framework, the first step is to identify candidate genes whose expression in cell type *c* is associated with the neighborhood cell type composition. To ensure compatibility across datasets and allow for comprehensive candidate gene selection, we augment the gene expression matrices **X**_1_ and **X**_2_ to 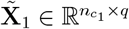 and 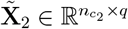, such that

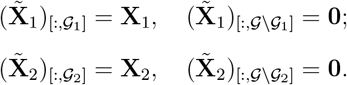

Similarly, we augment the neighborhood composition matrices **N**_1_ and **N**_2_ to 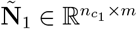 and 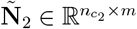, such that

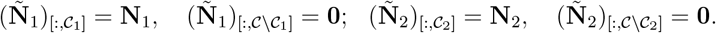

We then concatenate the augmented matrices to construct joint matrices:

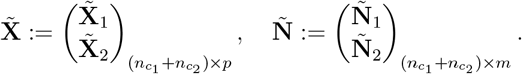

Following the procedure described in the single-condition case, we compute the covariance matrices of 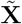 and 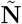, and select candidate genes via covariance tests. The significant genes are collected into a set 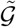, with size *q*^*′*^. We further denote the dataset-specific candidate gene sets as 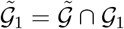 with size 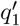 (dataset 1) and 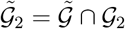 with size 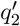 (dataset 2), and shared candidate gene set as 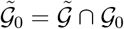 with size 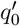.

To identify MCNs shared across the two datasets, we extract the expression matrix of shared candidate genes and the neighborhood composition matrix of shared cell types:

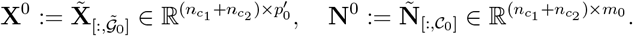

We then apply the nonnegative sparse CCA to **X**^0^ and **N**^0^ to identify *L*_0_ shared MCNs, yielding the pairs of gene and cell type coefficient vectors 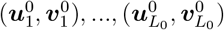.

After identifying the shared MCNs, we proceed to detect condition-specific MCNs. For illustration, we use dataset 1 as an example. In a manner analogous to the CCA procedure which iteratively adjusts for previously components, we subtract the share MCNs from **X**_1_ and **N**_1_ to obtain the residual matrices for dataset 1:

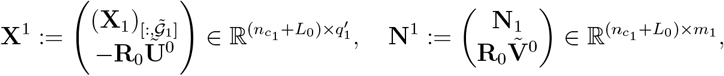

where

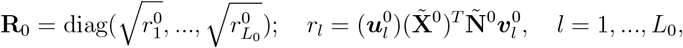

and 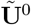 and 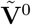 are row-stacked matrices of the augmented version of 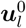 and 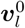:

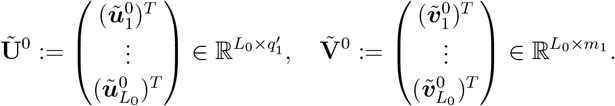

We then apply nonnegative sparse CCA to (**X**^1^, **N**^1^) to identify *L*_1_ dataset 1-specific MCNs, with the resulting pairs of gene and cell type coefficient vectors denoted as 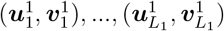. The same procedure is repeated for dataset 2 to obtain *L*_2_ dataset 2-specific MCNs, with pairs of gene and cell type coefficient vectors 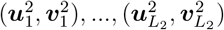.

### Parameter sensitivity and default settings

#### Sensitivity to kernel bandwidth *σ*

To assess the sensitivity of the covariance test to the Gaussian kernel bandwidth parameter *σ*, we performed a systematic analysis on the Xenium lymph node dataset using B cells and T cells as target cell types (Fig. S31). For each target cell type, we computed neighborhood cell type composition matrices with a series of *σ* values: 2, 5, 10, 20, 40, 60, 80, 100, 150, and 200. The neighborhood of each cell is weighted by a Gaussian kernel with bandwidth *σ*, followed by thresholding to zero out small weights. Thus, larger *σ* values should lead to broader neighborhoods involving more neighboring cells.

We first quantified the number of neighboring cells within a fixed radius (*r*_0_) around each target cell to assess the effective neighborhood size. As expected, the number of neighbors increased with *σ* for both B cells and T cells. At *σ* = 20, the median neighborhood size was 45 cells for B cells and 35 cells for T cells, with a well-distributed representation of diverse neighboring cell types.

We then performed covariance tests at each *σ* to identify genes whose expression in target cells was significantly associated with neighborhood composition. The number of significant genes (adjusted *p*-value *<* 0.05 or *<* 0.01) peaked when *σ* ranged between 10 and 40 for both B cells and T cells, while smaller or larger values of *σ* yielded fewer discoveries. Using *σ* = 20 as a baseline, we compared the overlap between significant gene sets across different *σ* values. For *σ* = 10 and *σ* = 40, the Jaccard index with the *σ* = 20 gene set was approximately 0.8, indicating substantial concordance. Most other *σ* values still yielded Jaccard indices above 0.6 (for B cells) or 0.5 (for T cells). Moreover, gene-level adjusted *p*-values from different *σ* settings were highly correlated with those at *σ* = 20, with Pearson correlation coefficients *R >* 0.95 across all comparisons (except for the extreme case of *σ* = 5).

These results demonstrate that the covariance test is robust to a wide range of *σ* values and that fixing *σ* (e.g., 20) is statistically sound and practically effective. For the Xenium lymph node dataset, the choice of *σ* = 20 strikes a balance between local resolution and statistical power: it defines neighborhoods of moderate size with adequate cellular diversity and yields the highest number of candidate genes for niche detection.

#### Default parameter settings

Unless otherwise specified, NicheScope analyses used the following default parameters: neighborhood cell type composition was computed using a Gaussian kernel with bandwidth *σ* = 20 and threshold *τ* = 0.05; genes associated with neighborhood composition were selected via a covariance test (*p <* 0.05), with a maximum of 500 candidate genes; and nonnegative sparse CCA was performed with penalty parameters *a* = 0.6 and *b* = 0.5.

### Quantification and statistical analysis

#### Neighborhood enrichment analysis of ST data

We constructed a spatial connectivity graph based on the Euclidean distances between cells, where each node represents a cell and an edge is drawn between two nodes if their spatial distance falls below a predefined threshold. Using this graph, we quantified the number of neighboring cell pairs for each pairwise combination of cell types. To assess whether specific cell type pairs co-localize more or less frequently than expected by chance, cell type labels were randomly permuted multiple times to generate a null distribution. A *z*-score was then calculated for each cell type pair, with a high positive or negative *z*-score indicating significant enriched or depleted co-localization, respectively. This analysis was performed using the functions `gr.spatial neighbors` and `gr.nhood_enrichment` from the *squidpy* Python package [103].

#### Clustering of neighborhood cell type composition

For a given target cell type *c*, we first compute its neighborhood cell type composition matrix using Eq. (1). The standardized version of this matrix is used to construct an AnnData object. We then perform Leiden clustering on the top principal components of the composition matrix at a specified resolution (e.g., 0.3) using the `tl.leiden` function from the *scanpy* Python package [104]. By summarizing the number of neighboring cells of each type within each cluster, we can infer which cell types are most commonly enriched in the neighborhood of target cells belonging to different clusters.

#### Pseudotime inference

Pseudotime inference of tumor cells in spatial transcriptomics data was performed using the *slingshot* R package [46]. Tumor cells were first extracted and used to construct a Seurat object, followed by data normalization using `SCTransform` and unsupervised clustering. Based on the resulting clusters, `slingshot` was applied to infer cell lineages and fit smooth lineage trajectories. Each cell’s pseudotime was then computed by projecting it onto the inferred lineage curves. It is important to note that pseudotime reflects the relative progression of a biological process as inferred from gene expression profiles, rather than representing actual chronological time.

#### Gene set enrichment analysis

To investigate the biological functions associated with a given MCN, we performed gene set enrichment analysis (GSEA) on niche genes with substantially nonzero coefficients, defined as *g*: ***u***_*g*_ *>* 0.05. Specifically, we employed an over-representation analysis (ORA) approach to assess whether the identified niche gene set was significantly enriched for known pathways or biological processes. Gene set libraries used in the analysis included MSigDB [105], Gene Ontology (GO) [106], KEGG [107], and Reactome [108]. Enrichment *p*-values were computed via the Enrichr web API [109], as implemented in the *gseapy* Python package [110]. Significantly enriched gene sets were then used to infer the key pathways and biological processes associated with the corresponding MCN.

#### Differentially expressed genes analysis

To gain a comprehensive understanding of a given MCN, we performed differential gene expression analysis for the corresponding niche cell types. For a specific niche cell type *k*, we first extracted the gene count matrix of all cells annotated as type *k* from the entire tissue section. These cells were then divided into two groups: within-MCN and outside-MCN. A cell was classified as within-MCN if it was located within a distance *r*_0_ of at least one target cell (of type *c*) with a positive niche score. Formally, the set of within-MCN cells of type *k* is defined as 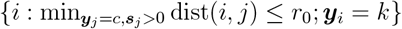. All remaining cells of type *k* were considered outside-MCN. DE genes between the within-MCN and outside-MCN groups were identified using the *DESeq2* R package [111], based on the count matrix and the two-group condition labels.

### Survival analysis

For the Xenium lung cancer dataset, we performed survival analysis to investigate whether the MCNs identified by NicheScope were associated with patient prognosis in lung cancer. Bulk RNA-seq data and clinical information for 507 LUAD patients were obtained from TCGA [57]. Clinical metadata included survival status (alive or deceased) and days to last follow-up. For living patients, the last follow-up time was treated as the survival time. For a given MCN of target cell type *c*, we selected a cell type *k* of interest (typically the target or one of the niche cell types) and divided cells of type *k* into two subgroups: a within-MCN subgroup *k*_1_ (referred to as MCN-like *k*) and an outside-MCN subgroup *k*_2_ (referred to as other *k*).

Using gene expression matrix extracted from the Xenium data containing subgroups *k*_1_, *k*_2_ of *k*, and the remaining *m* − 1 cell types as reference, we performed cell type deconvolution of the bulk RNA-seq profiles using RCTD [112]. This yielded patient-level cell type proportions 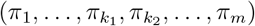. We then computed the relative abundance of the within-MCN subgroup *k*_1_ within the overall cell type *k* as 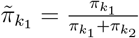. Patients were stratified into high and low groups based on the median of 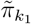, the estimated proportion of MCN-like cells of type *k*. We used the Kaplan-Meier estimator to estimate survival functions and compared them between the two groups. To quantify the prognostic association, we fitted a Cox proportional hazards model, in which the relative abundance of MCN-like *k* cells 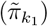 was used as a predictor, while adjusting for the overall proportion of cell type *k* as a covariate. The resulting hazard ratio (HR) reflects the relative risk of death for patients with a higher versus lower abundance of MCN-like *k* cells, compared to other *k* cells (i.e., subpopulation *k*_2_). A significantly elevated HR (i.e., HR *>* 1) indicates that the MCN-like subpopulation is associated with worse prognosis.

#### Ligand-receptor analysis

For the OpenST primary HNSCC and lymph node metastatic sections, which provide whole-transcriptome coverage, we employed the *liana* Python package [113] to perform ligand-receptor inference. For each MCN, we defined within-MCN cells as those located within a radius *r*_0_ of any target cell with a positive niche score: 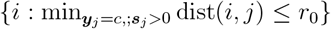. The gene expression matrix of these within-MCN cells was extracted and converted into an AnnData object. We used *liana*’s default ligand-receptor resource, a consensus network derived from literature-curated databases within OmniPath [114]. Inference was performed using `mt.rank aggregate`, which aggregates predictions from multiple ligand-receptor methods into consensus magnitude and specificity ranks. The magnitude rank reflects the overall interaction strength across methods, while the specificity rank quantifies how uniquely the interaction is associated with a particular cell type pair. Top-ranked ligand-receptor interactions, in terms of both magnitude and specificity, offered insights into the cell-cell communication landscape within each MCN.

### Data collection, preprocessing and analysis

We applied NicheScope to multiple spatial transcriptomics datasets generated by the Xenium and OpenST platforms, spanning normal and tumor tissues from lymph node, lung cancer, and HNSCC. Although the overall analysis pipeline remained consistent across datasets, we made dataset-specific adjustments to account for tissue context, cell type annotation confidence, and gene coverage. In all analyses, highly variable genes (HVGs) were selected to capture transcriptional heterogeneity within the target cell type. Unless otherwise noted, the top 3,000 HVGs were used. For the OpenST HNSCC dataset, 5,000 HVGs were selected to better capture the diversity across primary and metastatic tumors.

#### Xenium lymph node dataset

A formalin-fixed paraffin-embedded (FFPE) reactive lymph node sample was processed using the Xenium platform. The full dataset, comprising 708,983 cells and 5,001 genes (median 255 transcripts per cell), was downloaded from 10x Genomics. Two non-overlapping crops were extracted, each containing 197,644 and 192,378 cells, respectively. Gene expression counts were normalized to median library size and log1p-transformed. Cell type annotations were assigned by cell2location using a reference lymph node scRNA-seq dataset with 34 sub cell types [21]. Cells with annotation confidence (maximum normalized probability) *<* 0.3 were excluded, resulting in 80,534 and 74,851 cells for the two crops. Given the limitations of current spatial transcriptomics technologies, fine-grained sub cell type annotations can be less reliable due to transcriptional similarity and technical noise. We grouped these sub cell types into 13 major cell types and performed MCN detection at the major cell type level, thereby improving robustness and interpretability. For B cell-associated MCN detection, the analysis followed the NicheScope default pipeline. T cell-associated MCN detection followed the same protocol, with an additional filtering step to enhance signal robustness: genes expressed in fewer than 1% of T cells in the reference scRNA-seq were excluded prior to niche detection. This adjustment addressed the diffusely distributed nature of T cells in this dataset, which made niche signals more susceptible to noise from low-expression or misattributed transcripts. In contrast, B cells exhibited highly structured spatial organization, allowing the default pipeline to perform robustly without additional filtering.

### OpenST lymph node dataset

A normal lymph node tissue from a HNSCC patient was profiled using OpenST, yielding 82,518 cells and 24,663 genes. To assess the cross-platform reproducibility of NicheScope, only the 4,178 genes shared with the Xenium lymph node dataset were retained. After the same preprocessing and cell2location-based annotation, cells with confidence scores *<* 0.2 were excluded, resulting in 65,262 cells. Due to low confidence for T cell annotations, only B cell-associated MCN detection was performed using the default NicheScope parameters.

### Xenium lung cancer dataset

An FFPE lung adenocarcinoma was profiled using Xenium, yielding 278,328 cells and 5,001 genes (median 242 transcripts per cell). After normalization and log1p transformation, cell types were inferred via cell2location using a reference lung cancer scRNA-seq with 21 cell types [115], and cells with confidence scores *<* 0.5 were filtered out, resulting in 171,923 cells. Tumor cell, CD4^+^ T cell, and stromal cell-associated MCN detection were performed following the default NicheScope pipeline. In sparse CCA, the introduction of penalty terms may prevent earlier components from fully capturing weaker niche signals, which can instead emerge in later components. We indeed observed that some later components shared non-zero gene and cell type coefficients with earlier ones, and exhibited similar spatial distributions of niche scores and functions. Therefore, to more comprehensively represent such MCNs, spatially and functionally similar components were merged. The merged MCNs was defined by summing the gene and cell type coefficient vectors and normalizing them to unit *l*_2_ norm. In tumor cell-associated MCN detection, components 2 and 6 were merged into MCN2, and components 1, 5, and 9 into MCN3.

### OpenST HNSCC primary and metastatic tumor datasets

The primary and metastatic tumor samples were collected from a patient with HNSCC. For the metastatic tumor, we selected the deeply-sequenced section 28 from multiple available lymph node metastasis sections. The processed datasets contained 36,488 cells and 21,995 genes in the primary tumor, and 58,440 cells and 20,578 genes in the metastatic tumor. Gene expression counts for the two samples were normalized to the median library size and log1p-transformed separately. Cell type annotations for the primary tumor combined the original labels with those transferred by cell2location using a reference HNSCC scRNA-seq [116], whereas the metastatic tumor used the original annotation directly. In both datasets, subtypes were merged and relabeled to generate a unified annotation of 12 major cell types. Shared tumor cell niches were identified by concatenating the datasets and applying NicheScope with a Gaussian kernel bandwidth of *σ* = 40.For condition-specific niche discovery, the first 8 shared CCA components were adjusted before applying NicheScope separately on the primary and metastatic tumors using the same parameters.

## Data availability

Xenium spatial transcriptomics datasets, including the reactive lymph node and lung adenocarcinoma samples, were obtained from 10x Genomics: the lymph node dataset (https://www.10xgenomics.com/cn/datasets/preview-data-xenium-prime-gene-expression) and the lung cancer dataset (https://www.10xgenomics.com/cn/datasets/xenium-human-lung-cancer-post-xenium-technote). OpenST datasets, comprising healthy lymph node, primary HNSCC, and lymph node metastasis samples, were downloaded from GEO under accession num-ber GSE251926. Reference scRNA-seq datasets used for cell type annotation were obtained from publicly available sources: lymph node (Zenodo, https://zenodo.org/records/3710410), lung cancer (CellxGene, https://cellxgene.cziscience.com/collections/edb893ee-4066-4128-9aec-5eb2b03f8287), and primary HNSCC (GEO, GSE103322). TCGA-LUAD bulk RNA-seq and clinical data were accessed via the GDC portal (https://portal.gdc.cancer.gov/projects/tcga-luad).

## Code availability

The IBSEP software and scripts for reproducing the analyses are available at: https://github.com/xinyiyu/NicheScope.

